# Otoprotective effect of MnTBAP in cisplatin-induced hearing loss

**DOI:** 10.64898/2026.06.04.730129

**Authors:** Shomaila Mehmood, Pankaj Bhatia, Samson Jamesdaniel

## Abstract

**Objective:** Cisplatin, a life-saving chemotherapeutic drug, causes ototoxicity. Although sodium thiosulfate is used to prevent ototoxicity in pediatric patients, no other intervention has been approved for clinical use against cisplatin-induced hearing loss. Hence, there is an urgent need to identify drugs that prevent cisplatin ototoxicity.

**Methods:** CBA/J mice were treated with cisplatin (3 mg/kg, i.p., daily for 5 days), and MnTBAP (10 mg/kg, i.p., daily for 8 days) was used to inhibit cisplatin-induced ototoxicity. Auditory brainstem responses (ABRs) and distortion product otoacoustic emissions (DPOAEs) were recorded before and after treatment to assess hearing loss, while immunohistochemistry was performed to examine hair cells and spiral ganglion neuron (SGN) loss.

**Results:** Cisplatin treatment elevated the nitrotyrosine levels in hair cells and SGNs and increased the loss of these cells in the middle and basal cochlear regions. A negative correlation was observed between cisplatin-induced changes in the hair cell count or SGN density and nitrotyrosine levels. Cisplatin elevated the hearing thresholds and lowered the DPOAE amplitudes. However, MnTBAP cotreatment prevented the cisplatin-induced changes in the hearing sensitivity and reversed the morphological changes.

**Conclusion:** The otoprotection observed with MnTBAP cotreatment indicates its potential as a therapeutic drug against cisplatin-induced ototoxicity.

## Introduction

Cisplatin is a highly used, efficient anti-cancer treatment against a wide range of solid tumors [1–4]. However, it has significant adverse effects, some of which might impair a patient’s quality of life permanently after treatment. Particularly, cisplatin treatment causes hearing loss in approximately 60% of patients, including both pediatric and adult cancer populations, which affects their quality of life [5, 6, 7]. Both children and adults develop high-frequency hearing loss after cisplatin treatment, which can progress to cause considerable damage in frequency regions essential to understand human language [8–10]. Cisplatin-induced ototoxicity can result in mild to severe hearing impairment and is influenced by age, concurrent cranial irradiation, pre-existing hearing loss, dosing schedules, and cumulative dose [11, 12]. Studies have indicated that the three primary targets of cisplatin in the cochlea are the sensory hair cells in the organ of Corti [13], the lateral wall tissues [14], and the spiral ganglion neurons (SGNs) in the modiolus [15]. Cisplatin treatment causes the death of outer hair cells (OHCs) in the basal turn of the cochlea; therefore, it affects high-frequency hearing more than low-frequency hearing [16, 17]. Though the cytotoxic mechanism underlying cisplatin-induced ototoxicity is not fully understood, various studies have reported that cisplatin-induced cochlear apoptosis is intricately linked to oxidative stress, DNA damage, and inflammatory factors [11, 18–20]. Moreover, among the cochlear cytotoxic mechanisms, oxidative stress plays a major role, as cisplatin activates the NOX3 enzyme, increasing the generation of superoxide radicals in the inner ear [11, 21]. Additionally, the iNOS pathway is activated, leading to nitric oxide (NO) generation during cisplatin-induced ototoxicity [22, 23]. The reaction of NO with superoxide radicals produces peroxynitrite (ONOO^-^) that ultimately causes protein nitration. Nitrated proteins can alter the signaling pathways that regulate apoptosis and cause damage to hair cells in the cochlea [24]. Nitration of LMO4, a transcriptional regulator involved in cell survival/death and inner ear development [25–29], has been implicated as a key mediator of cisplatin ototoxicity [24]. Thus, nitrative stress has emerged as a critical factor in cisplatin-induced hearing loss.

To mitigate cisplatin ototoxicity, numerous therapeutic strategies have been employed, among which reducing the pro-inflammatory cytokine release has shown efficacy [30–33]. Other approaches, such as the use of antioxidants sodium thiosulfate (STS) and N-acetylcysteine (NAC), reduce the intracellular reactive oxygen species (ROS) accumulation to attenuate cisplatin-induced hearing loss [34–38]. However, STS and NAC can interfere with the anti-cancer activity of cisplatin [39, 40] and compromise the chemoprotection [40, 41]. Therefore, it is important to identify novel interventional drugs that are highly effective in mitigating cisplatin-induced ototoxicity without interfering with its anticancer efficacy [31, 42–44].

Nitrative stress has emerged as a promising target for preventing cisplatin ototoxicity. Metalloporphyrins such as FeTMPyP and MnTBAP chloride (Mn (III)tetrakis (4-benzoic acid) porphyrin chloride) are efficient ONOO^-^ scavengers [45], which probably contributes to their efficacy in attenuating cisplatin-induced apoptosis, nitrative stress, and nephrotoxicity [46]. They also prevent meningitis-related hearing loss resulting from increased reactive oxygen species (ROS) and ONOO^-^ [47]. Other PNDCs, such as SRI110, a Mn (III) complex, have been reported to prevent cisplatin-induced cytotoxicity and nitrative stress [48]. MnTBAP is widely used *in vivo* models of oxidative stress injuries and has been reported to selectively target ONOO^-^ when it is in a pure form [45]. As MnTBAP has been reported to inhibit both superoxide and ONOO^-^ and targeting of ONOO^-^ by metalloporphyrins is emerging as an efficient interventional strategy to mitigate cisplatin ototoxicity, we assessed the efficacy of MnTBAP cotreatment against cisplatin ototoxicity by evaluating the effect of MnTBAP cotreatment on hair cells, SGNs, and auditory function using a mouse model.

## Materials and methods

### Animals

A total of 12 male and 12 female CBA/J mice (Six-week-old), obtained from Jackson Labs (Jackson Laboratories, Bar Harbor, ME, USA), were kept in a temperature-controlled room with 12-hour light/dark cycles in the laboratory animal facility. All animal procedures received approval from the Institutional Animal Care and Use Committee (IACUC) at Wayne State University (IACUC-23-07-5961), and experiments were conducted in accordance with applicable guidelines, regulations, and ARRIVE reporting standards. All animals were allowed a week to acclimatize and were carefully monitored for changes in overall health and activity that may have resulted from cisplatin treatment.

### Drug administration

The animals were treated with cisplatin following an 8-day protocol (Fig. 1). Mice were assigned to one of four experimental groups: control, cisplatin, MnTBAP+Cisplatin, and MnTBAP alone, maintaining a balance of males and females in each group. Animals in the cisplatin treatment group received a gradual intraperitoneal (i.p.) infusion of veterinary-grade cisplatin (Accord Healthcare, NDC16729-288-38, USA) diluted in sterile saline (pH 7.4) at a dose of 3 mg/kg body weight each day for 5 days while the animals in the MnTBAP cotreatment group received i.p. injections of 10 mg/kg of MnTBAP (Catalog no. sc-221954A, Santa Cruz Biotechnology) each day. MnTBAP was dissolved in sterile saline and administered 3 hours before cisplatin injections for 5 days, followed by MnTBAP treatment alone for the remaining 3 days. The MnTBAP alone group received MnTBAP on all 8 days, while an equivalent volume of saline was administered to the control animals. All animals were subcutaneously hydrated with 1 mL of 0.9 % saline every day.

**Fig. 1.**
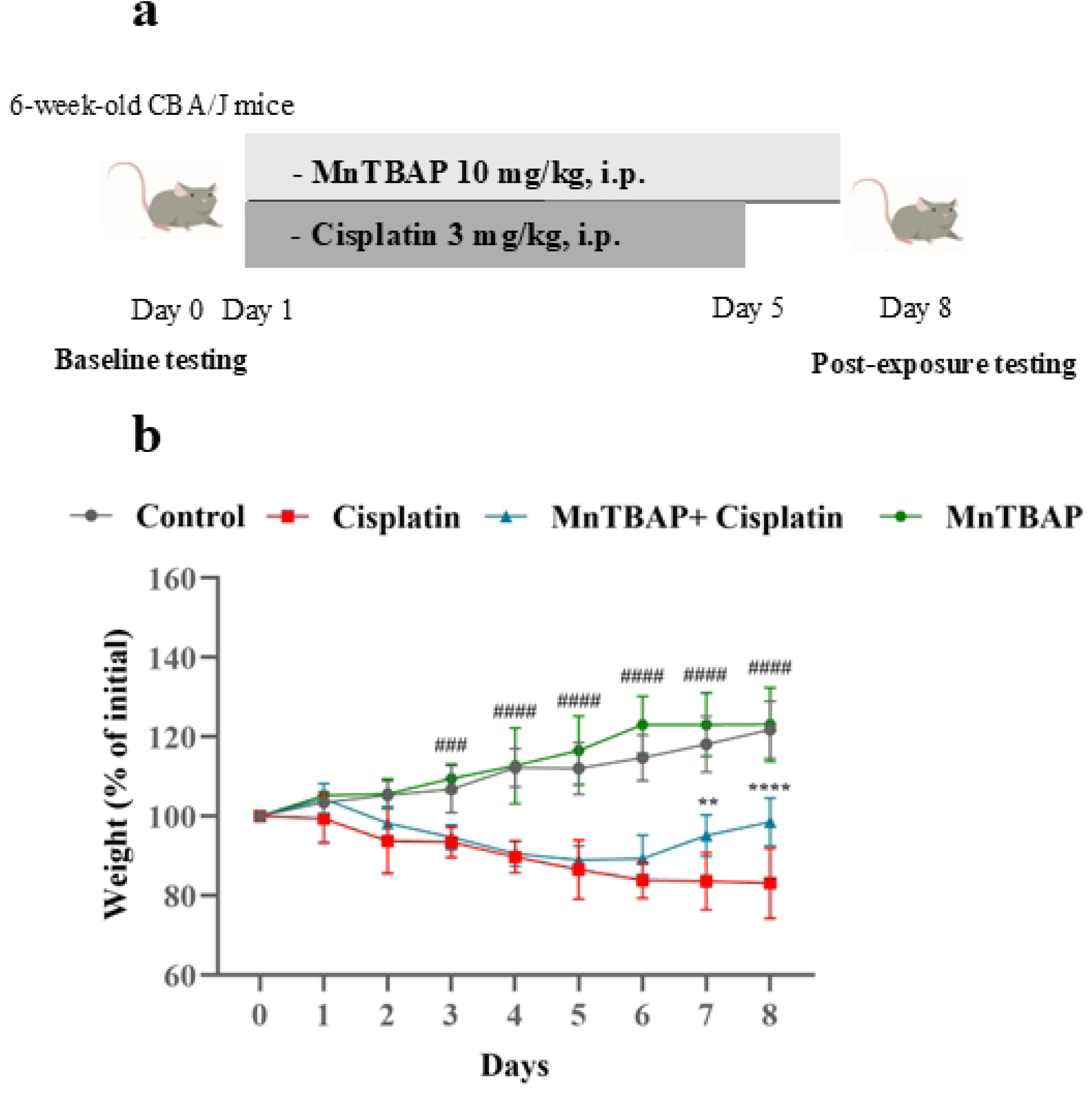
Experimental protocol and weight monitoring. **(a)** Cisplatin was administered for 5 days (3 mg/kg, i.p.), MnTBAP cotreatment (10 mg/kg, i.p.) was given for 5 days, 3 hours before cisplatin injection, followed by MnTBAP alone for 3 days. MnTBAP alone group received MnTBAP at the same dosage. Auditory testing was performed before the treatment began and on day 8 after the completion of the treatment protocol. Mice were given 1 mL sterile saline every day for 8 days to avoid dehydration. **(b)** Weight change over the 8 days was monitored for animals in all groups. MnTBAP cotreatment prevented cisplatin-induced weight loss. ** indicates p < 0.01, ****p < 0.0001, MnTBAP+Cisplatin compared to cisplatin alone; ^###^ indicates p < 0.001, ^####^p < 0.001, cisplatin relative to control. Control animals were treated with a saline solution. Data are expressed as mean±S.D. (n=5-6).

### Hearing tests

Hearing was tested in all mice by recording auditory brainstem responses (ABR) and distortion product otoacoustic emissions (DPOAEs), before and after drug administration. Auditory testing was conducted using Tucker Davis Technologies (TDT) System III hardware, and sound stimuli were presented using the BioSigRZ software (v5.7.6.). Mice were put on a temperature-controlled heating pad kept at 38 °C within the sound-proof chamber (ECKEL-AB-4230, Morrisburg, Ontario) after being anesthetized with isoflurane (4 % induction, 1.5 % maintenance with 1 L/min O_2_). For ABR recordings, broadband click stimuli (90-20 dB SPL, 5 dB intervals) and tone bursts at 4, 8, 16, 24, and 32 kHz frequencies (90-25 dB SPL, 5 dB intervals) were delivered. To obtain the ABR waveforms, 512 stimulus presentations delivered at 21 Hz were averaged. The hearing threshold was recorded as the lowest stimulation level at which a waveform with a discernible peak was observed.

Two primary tones, f1 and f2, were used to assess DPOAEs at a f2/f1 ratio of 1.2. For L1 levels ranging from 80 dB SPL to 20 dB SPL in 10 dB increments, the L2–L1 was maintained at +10 dB. The stimulus was produced using the Tucker-Davis Technology (TDT) RZ6 system, and the multifield magnetic speakers (TDT, Alachua, FL, USA) were used to provide f1 and f2. The f2 frequency ranged between 4 and 40 kHz. The ER10B+probe microphone (Etymotic Research, Inc., Elk Grove Village, IL, USA) and TDT hardware and software were used to measure SPLs at the cubic difference frequency (2f1–f2). Distortion product data were collected every 20.971 ms and averaged 512 times, while a noise floor was measured using a 100 kHz band around 2f1–f2.

### Immunohistochemistry

Following euthanasia, the temporal bones were promptly removed in ice-cold PBS, and the blood in the cochlea was flushed out by perfusing it with 4 % paraformaldehyde (PFA) in 1x PBS (pH 7.4) through the oval window. The cochlea was decalcified using 100 mM ethylenediaminetetraacetic acid disodium salt (EDTA) solution in 1x PBS (pH 7.4) for almost 48-72 hours. After decalcification, the tissues were microdissected under a stereomicroscope (Leica model # M 165 C) in ice-cold PBS. The cochlear sensory epithelium was separated from the lateral wall and bone capsule and was divided into apex, middle, and basal parts. Each cochlear section was initially blocked with 10 % normal goat serum (NGS) for 1 hour at room temperature and washed thrice with 1x PBS (pH 7.4). Following this, the cochleae were incubated in two sets of primary and secondary antibodies. The first set included mouse monoclonal Myosin-VIIa (Santa Cruz Biotechnology, catalog no. sc-74516, 1:500) and goat anti-mouse Alexa Fluor^TM^ 488 (Thermo Fisher Scientific, catalog no. 21131, 1:500) while the second set consisted of mouse monoclonal nitrotyrosine (Santa Cruz Biotechnology catalog no. sc-32757, 1:400) and goat anti-mouse Alexa Fluor^TM^ 568 (Thermo Fisher Scientific, catalog no. 21124, 1:500). The tissue sections were incubated in the primary antibodies overnight at 4 °C followed by incubation with secondary antibodies for 2 hours at room temperature. The specimen were further incubated with Alexa Fluor^TM^ 488 Phalloidin (Invitrogen^TM^, Thermo Fischer Scientific, Catalog no. A12379, 1:40) for 30 minutes at room temperature and then mounted with ProLong^TM^ Gold antifade with DAPI (Invitrogen^TM^, Thermo Fischer Scientific, catalog no. P36935). The Biotium CoverGrip^TM^ Coverslip Sealant (Thermo Fisher Scientific, catalog no. NC0154994) was used to seal the coverslips.

### Confocal microscopy and image analyses

High-resolution confocal images were collected with Carl Zeiss Laser Scanning Systems (Zeiss LSM-800) using an oil-immersion 40× objective. The hair cells in the control and drug-treated groups were counted using myosin-VIIa staining in 50 µm cochlear sections [49]. The fluorescence intensity of nitrotyrosine staining was measured in 12-14 OHCs [50] and 3 SGNs per image. The SGN density per 10^4^ µm^2^ was calculated following the protocol used by Hu et al. (2021) using ImageJ/Fiji software (v 1.46r) [51].

### Statistical analyses

All the experiments were performed in 5-6 biological replicates. A two-way analysis of variance (ANOVA) test followed by Tukey’s multiple comparisons test was used for statistical analyses, and a p-value of < 0.05 was considered statistically significant. The association between cisplatin-induced changes in nitrotyrosine levels and hair cell count and SGN density was determined using Pearson’s correlation coefficient at 95 % confidence interval. All data were analyzed using GraphPad Prism software (v10.2.3, La Jolla, CA, USA) and expressed as mean ± standard deviation (S.D.) or mean ± standard error mean (S.E.M).

## Results

### MnTBAP cotreatment prevented cisplatin-induced weight loss

Body weight was monitored daily throughout the 8-day treatment protocol (Fig. 1a) to assess the systemic effects of cisplatin and the potential protective role of MnTBAP. Mice receiving cisplatin alone exhibited a progressive decline in body weight starting from day 2, reaching a significant reduction on day 8 compared to control animals. In contrast, mice cotreated with MnTBAP exhibited significantly less weight loss, with body weight on day 8 remaining substantially higher than that of the cisplatin-treated group. The MnTBAP alone group maintained body weights comparable to controls throughout the study period, indicating MnTBAP treatment alone did not alter the body weight of animals (Fig. 1b).

### Cisplatin treatment increased nitrotyrosine levels in hair cells and SGNs

Immunohistochemistry of the apex, middle and basal cochlear turns with anti-nitrotyrosine indicated that cisplatin treatment increased the nitrotyrosine levels in OHCs (Fig. 2a). The quantification of nitrotyrosine staining intensity indicated that cisplatin caused a significant increase in nitrotyrosine levels across all the three regions of the cochlea as compared to the control (Fig. 2b). Likewise, cisplatin treatment also increased the nitrotyrosine levels in the SGNs in the middle and basal cochlear turns, compared to the controls (Fig. 2c and d). These results indicate that cisplatin treatment induces nitrative stress in the cochlea.

**Fig. 2.**
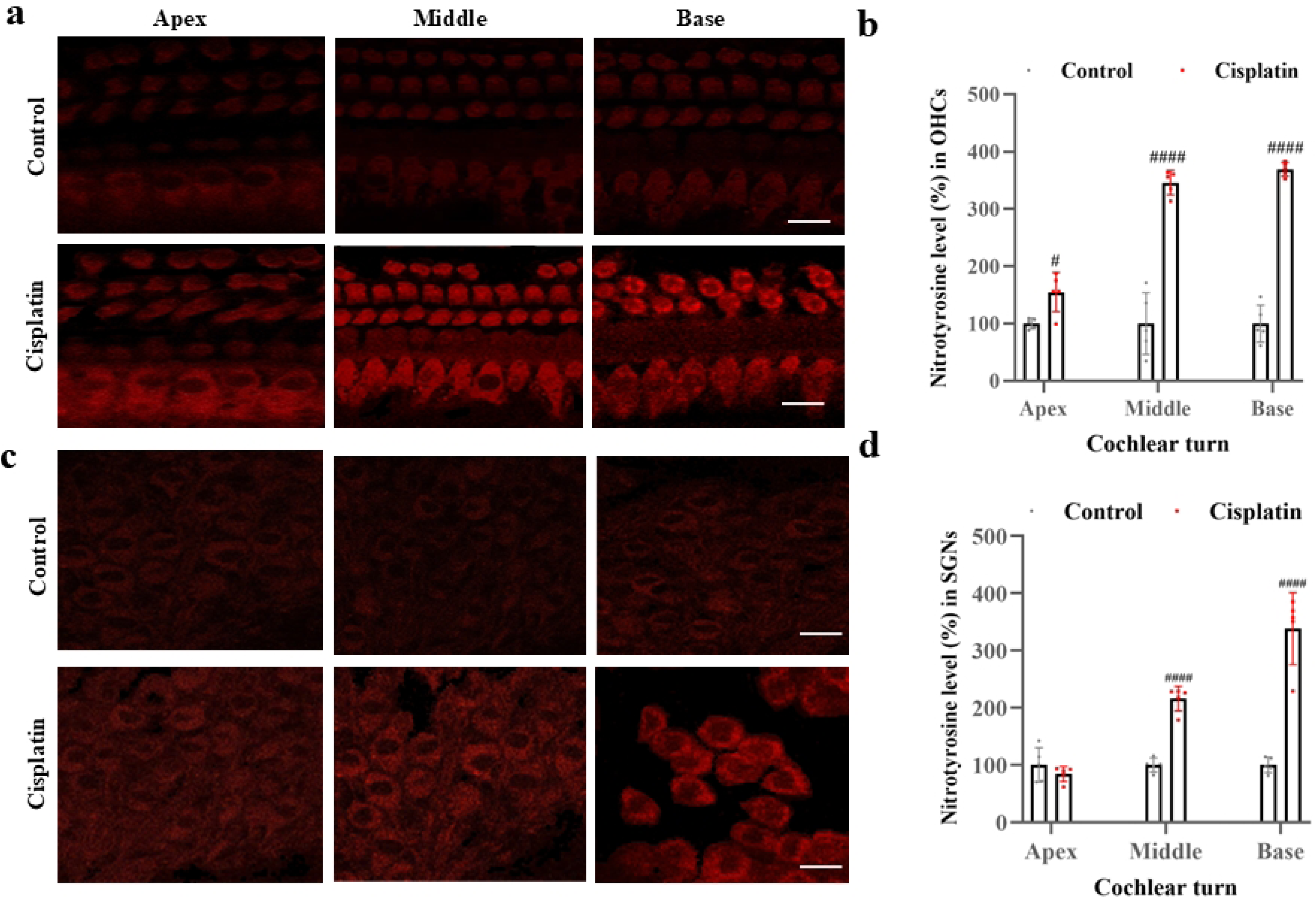
Nitrotyrosine levels in hair cells and SGNs. **(a)** Immunohistochemistry with nitrotyrosine antibody showed that cisplatin treatment increased nitrotyrosine levels (red) in OHCs in the apex, middle, and basal turns as compared to controls. **(b)** Quantification of nitrotyrosine staining intensity indicated that cisplatin significantly increased the nitrotyrosine levels across all three cochlear regions. **(c)** Cisplatin treatment increased nitrotyrosine levels in the SGNs in the apex, middle, and basal turns of the cochlea. **(d)** Quantification of nitrotyrosine staining intensity indicated a significant increase in nitrotyrosine levels in the middle and basal SGNs in the cisplatin-treated cochlea. ^#^ indicates p < 0.05, ^####^p < 0.0001, cisplatin relative to control. Saline-treated mice were used as controls. Data are expressed as mean±S.D. (n=5). Scale bar = 20 μm. OHCs, outer hair cells; SGNs, spiral ganglion neurons.

### MnTBAP cotreatment mitigated cisplatin-induced OHC loss

Immunohistochemical analysis with anti-Myosin-VIIa was used to assess the hair cell loss in three cochlear regions across all four groups (Fig. 3a). This analysis indicated that cisplatin treatment resulted in OHC loss, particularly in the middle and basal regions, as compared to the control and MnTBAP alone groups. There was no loss of inner hair cells (IHCs) in all groups. However, MnTBAP cotreatment prevented the cisplatin-induced loss of OHCs in the middle and basal turns. The count of IHCs and OHCs (Fig. 3b and c) indicated that MnTBAP cotreatment mitigates cisplatin ototoxicity by preventing the death of OHCs after cisplatin treatment.

**Fig. 3.**
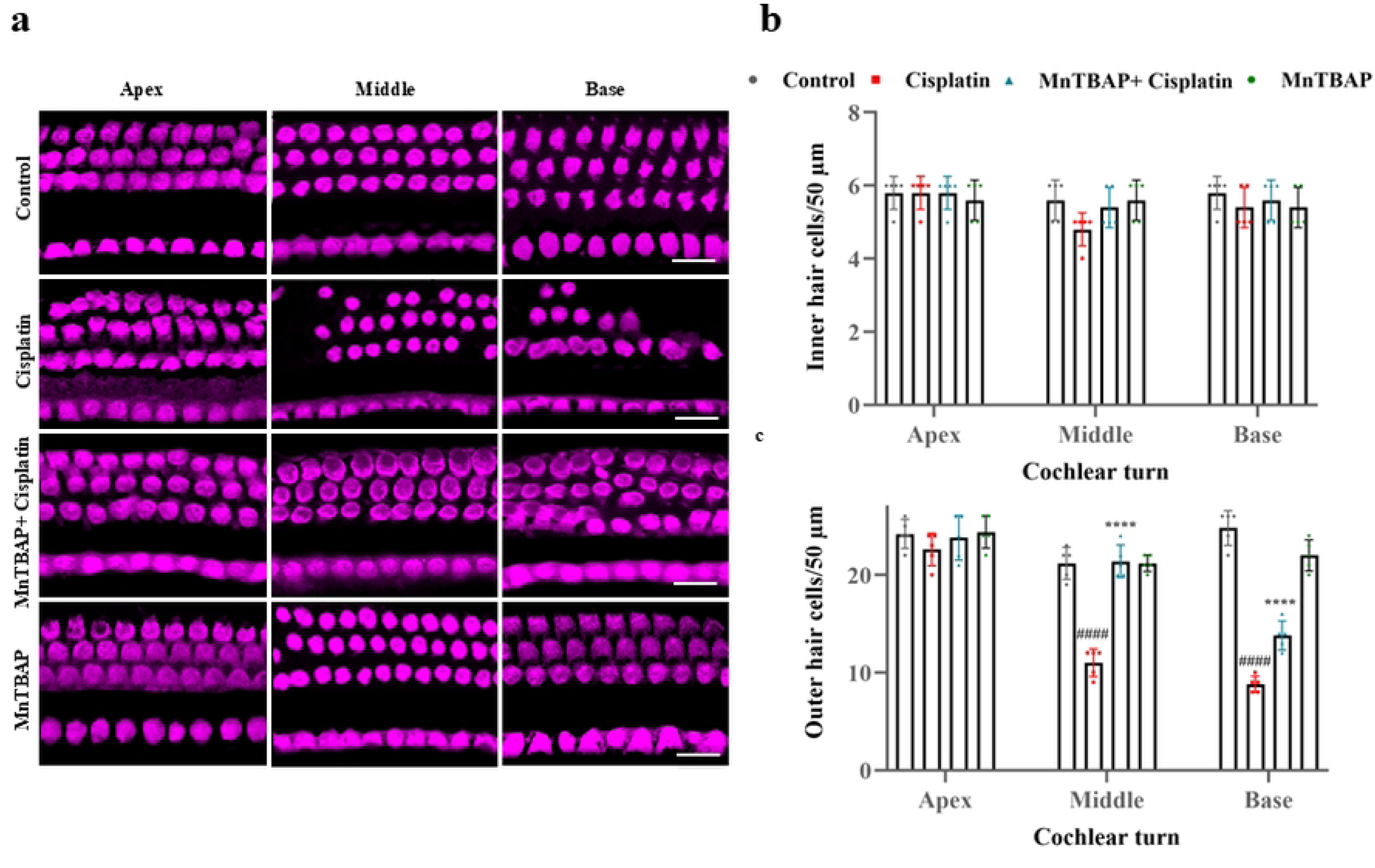
Effect of MnTBAP cotreatment on cisplatin-induced OHC loss. **(a)** Immunohistochemistry with anti-Myosin-VIIa indicated a significant loss of OHCs in cisplatin-treated mice in the middle and basal turns of the cochlea compared with controls. However, MnTBAP cotreatment significantly mitigated cisplatin-induced OHCs loss. **(b)** and **(c)** show the numbers of IHCs and OHCs in all three cochlear turns. These graphs illustrate significant loss of OHCs in cisplatin-treated mice in the middle and basal regions and no loss of IHCs. **** indicates p < 0.0001, MnTBAP+Cisplatin compared to cisplatin alone. ^####^ indicates p < 0.0001, cisplatin relative to control. Saline-treated mice were used as controls. Data are expressed as mean±S.D. (n=5). Scale bar = 10 µm. IHC, inner hair cells; OHCs, outer hair cells.

### MnTBAP cotreatment mitigated cisplatin-induced SGN loss

The SGNs were visualized by staining actin with phalloidin, which indicated that cisplatin treatment significantly reduced the SGN density in the middle and basal regions of the cochlea. However, MnTBAP cotreatment significantly attenuated the cisplatin-induced decrease in the SGNs density in the middle and basal cochlear regions (Fig. 4a). The quantification of SGN density revealed a higher SGN loss in the middle and basal turns in the cisplatin-treated cochlea, which was significantly reversed by the MnTBAP cotreatment, particularly in the basal turn (Fig. 4b). This indicates that MnTBAP prevents cisplatin-induced SGN loss in the cochlea.

**Fig. 4.**
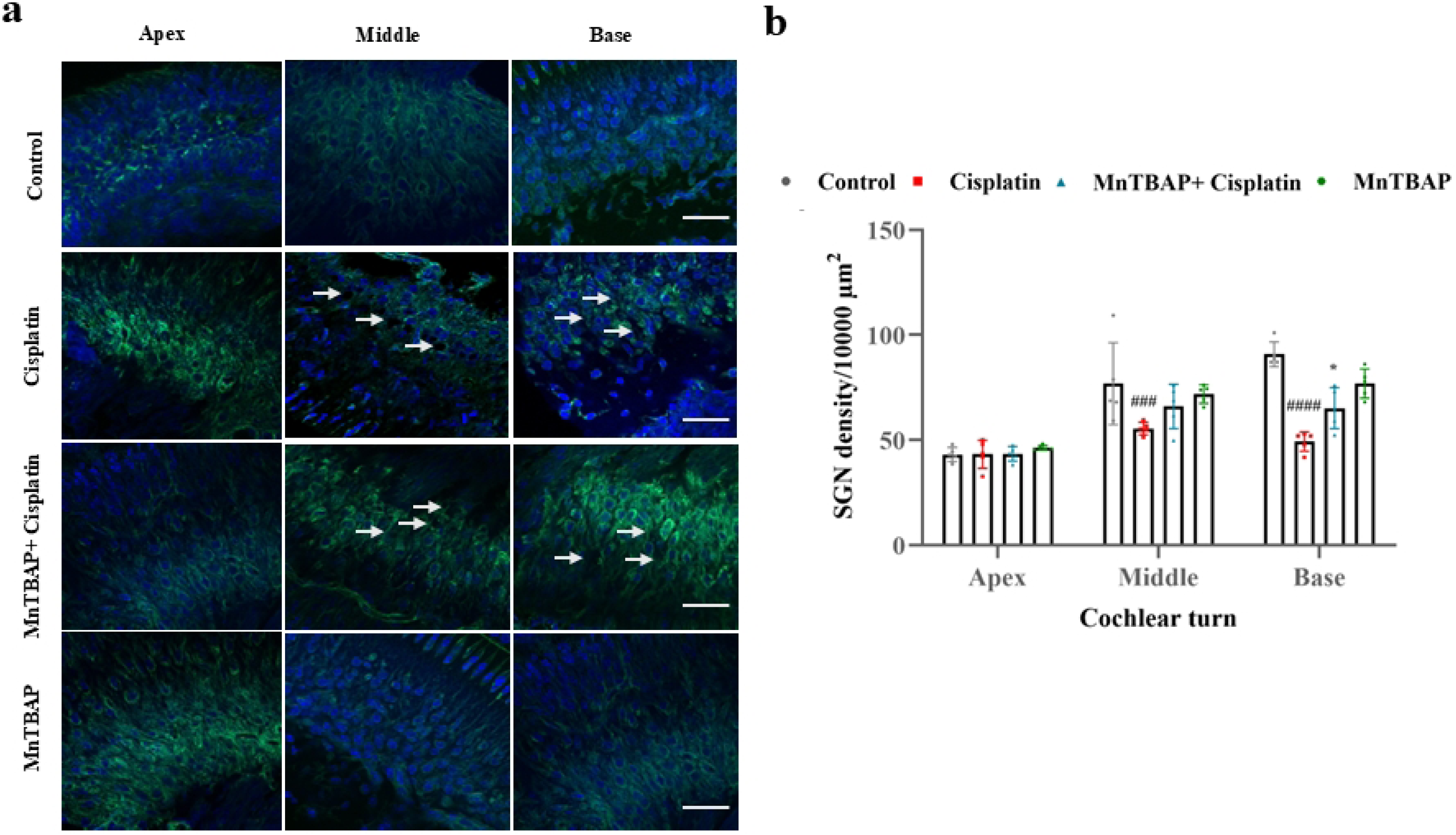
Effect of MnTBAP cotreatment on cisplatin-induced SGN loss **(a)** Cisplatin treatment resulted in the loss of SGNs in the middle and basal turns of the cochlea, as shown by the phalloidin (green) and DAPI (blue) staining. However, MnTBAP cotreatment prevented the cisplatin-induced SGN loss in the middle and basal turns of the cochlea. **(b)** The quantification of SGN density indicated that cisplatin treatment significantly decreased the SGN density, which was reversed by MnTBAP cotreatment. *indicates p < 0.05, MnTBAP+Cisplatin compared to cisplatin alone. ^###^ indicates p < 0.001, ^####^p < 0.0001, cisplatin relative to control. Saline-treated mice were used as controls. Data are expressed as mean±S.D. (n=5). Scale bar = 20 µm. SGNs, spiral ganglion neurons.

### Cisplatin-induced changes in nitrotyrosine level correlated with OHC and SGN loss

The correlation analysis between cisplatin-induced changes in nitrotyrosine levels and hair cell count using simple linear regression model indicated a strong negative correlation between the two variables (R^2^ = 0.9353, Fig. 5a). Similarly, nitrotyrosine levels and SGNs density also revealed a negative correlation (R^2^ = 0.9069, Fig. 5b). This further validates the fact that higher nitrotyrosine levels after cisplatin treatment probably contributes to OHC and SGN death in the cochlea.

**Fig. 5.**
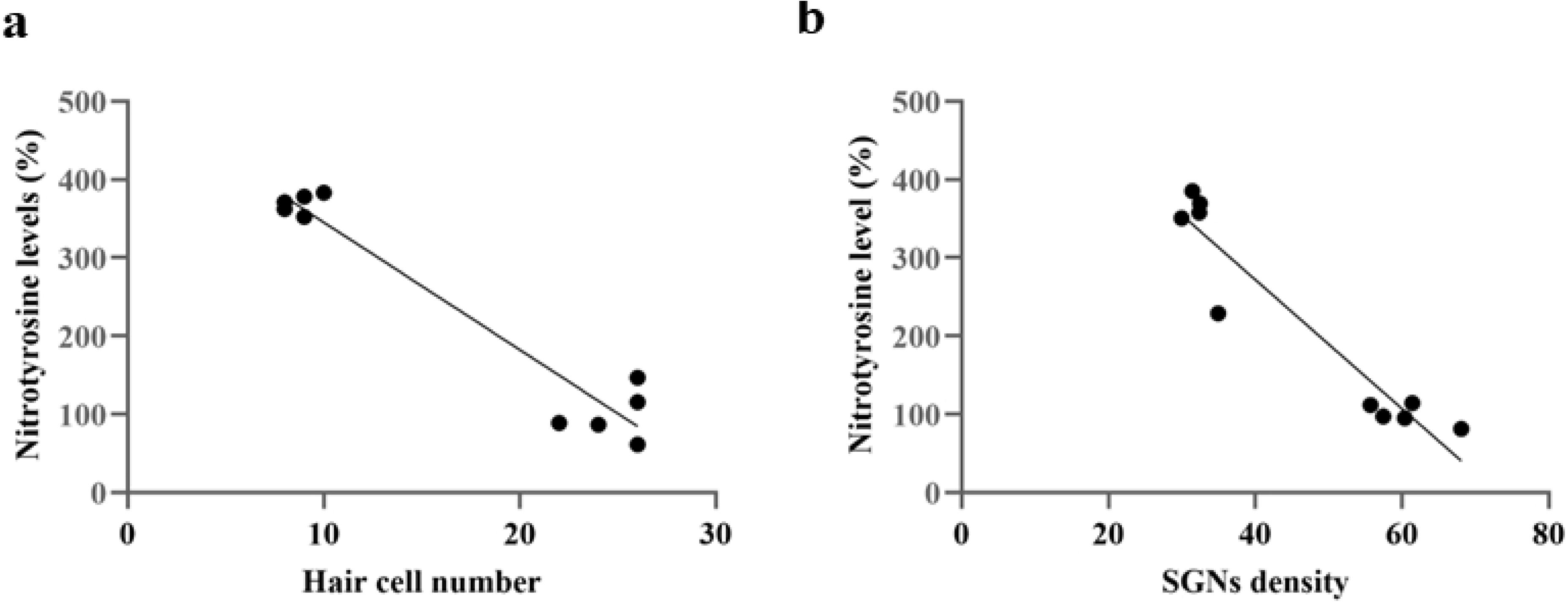
Correlation between nitrotyrosine levels and cisplatin-induced hair cell and SGN loss. Cisplatin-induced increase in nitrotyrosine levels showed a strong correlation with changes in hair cell count and SGN density. **(a)** and **(b)** Analysis of correlation between cisplatin-induced changes in the hair cell count, SGNs density, and nitrotyrosine levels indicated a negative correlation (R^2^ = 0.9353 and R^2^ = 0.9069, respectively with p < 0.001. (n=5). SGNs, spiral ganglion neurons.

### MnTBAP cotreatment attenuated cisplatin-induced hearing loss

Auditory sensitivity was assessed using ABR recordings after cisplatin exposure, which indicated that cisplatin treatment induced a significant increase in the ABR thresholds with threshold shifts ranging from 9 to 14 dB. However, cotreatment with MnTBAP, attenuated the cisplatin-induced increase in ABR thresholds and threshold shifts (Fig. 6a, b). Similarly, cochlear OHC function was assessed using DPOAEs, which showed a significant decrease in DPOAE amplitudes in cisplatin-treated mice (Fig. 6c). The DPOAE amplitudes were significantly lower in the mid-high frequency regions (16 and 24 kHz). However, MnTBAP cotreatment reversed the cisplatin-induced decrease in the DPOAE amplitudes. This indicates that MnTBAP is protective against cisplatin-induced changes in hearing sensitivity and restores the OHC activity.

**Fig. 6.**
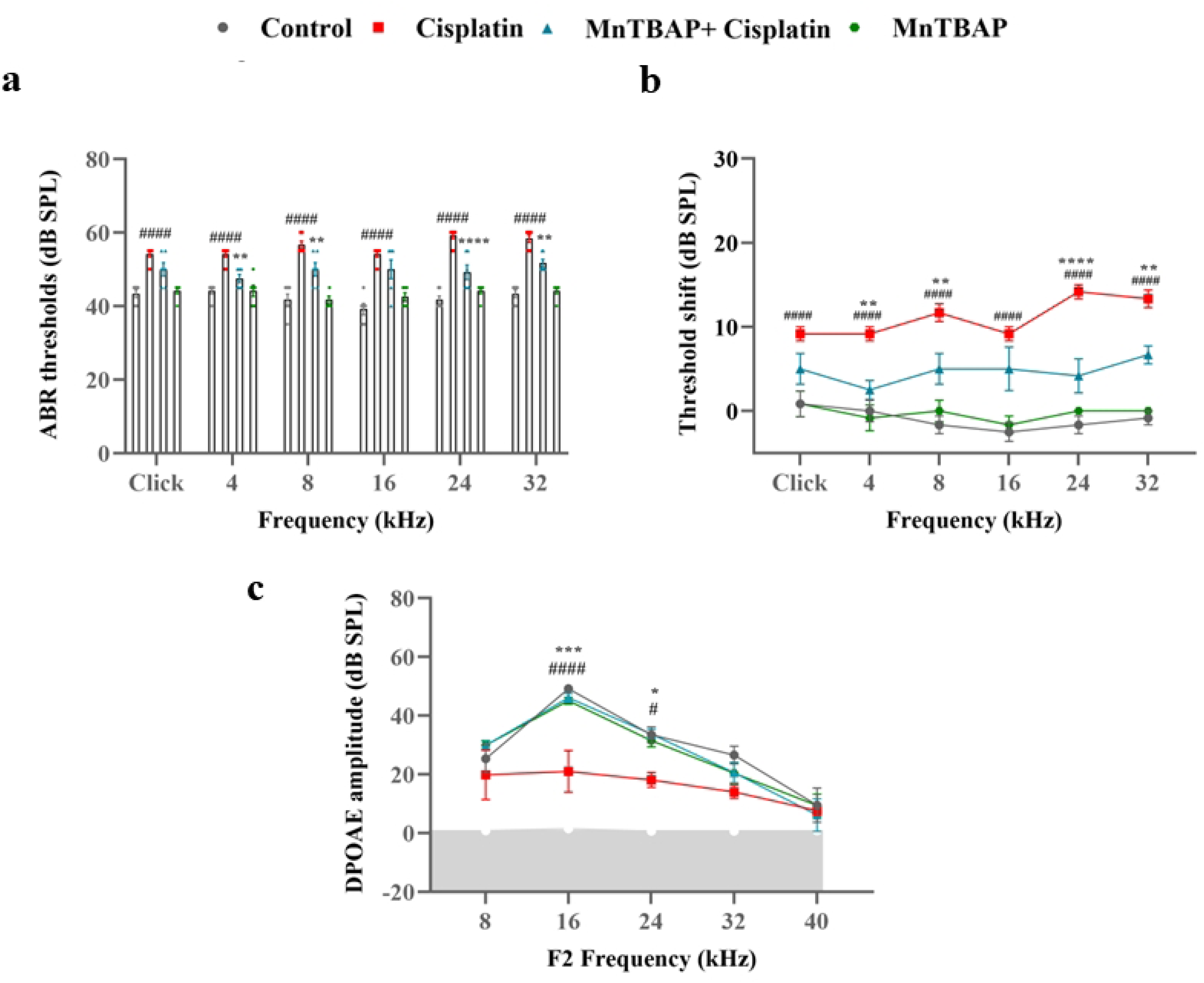
Effect of MnTBAP cotreatment on cisplatin-induced hearing loss. **(a)** The ABRs indicated that cisplatin-treated mice showed significantly higher hearing thresholds than control mice at multiple frequencies. However, MnTBAP cotreatment significantly attenuated the cisplatin-induced increase in hearing thresholds at 4, 8, 24, and 32 kHz. **(b)** Cisplatin-treated mice showed substantially higher threshold shifts than control mice; however, MnTBAP cotreatment significantly attenuated the cisplatin-induced hearing threshold shifts. **(c)** The DPOAEs recorded using 8, 16, 24, 32, and 40 kHz f2 stimuli indicated that the DPOAE amplitudes elicited using 70 dB SPL (L1) stimuli were significantly decreased in cisplatin-treated mice at 16 and 24 kHz frequencies. However, MnTBAP cotreatment significantly attenuated the cisplatin-induced decrease in DPOAEs amplitude at 16 and 24 kHz. Gray shaded region indicates the mean biological noise floor. Data recorded from the left ear is illustrated. * indicates p < 0.05, **p < 0.01, ***p < 0.001, ****p < 0.0001, MnTBAP+Cisplatin compared to cisplatin alone. ^#^ indicates p < 0.05, ^####^p < 0.0001, cisplatin relative to control. Saline-treated mice were used as controls. Data are expressed as mean±S.E.M. (n=5-6). ABR, auditory brainstem response; DPOAE, distortion product otoacoustic emissions.

## Discussion

Cisplatin is widely used to treat numerous types of cancers, which include nasopharyngeal cancer, lung cancer, malignant lymphoma, and other tumors [4]. However, nephrotoxicity, neurotoxicity, and ototoxicity are the side effects of cisplatin, which limit its usage. Amongst these major side effects, ototoxicity manifests as irreversible, progressive neurosensory hearing loss [11]. Hence, it is crucial to develop preventive measures to mitigate cisplatin-induced ototoxicity. The present study indicated that nitrotyrosine levels are elevated in OHCs and SGNs, and cisplatin treatment results in increased hair cell and SGN loss. A strong association between high nitrotyrosine levels and hair cell and SGN loss in the cochlea was observed after cisplatin exposure. More importantly, MnTBAP cotreatment protected hair cells and SGNs from cisplatin-induced death and prevented cisplatin-induced hearing loss.

Oxidative stress plays a crucial role in cisplatin-induced cochlear cell death [11], as NOX3 enzyme is activated and superoxide radicals are produced in the inner ear. Additionally, cisplatin treatment activates the iNOS pathway, resulting in the production of NO in the cochlea [22, 23]. The NO reacts with superoxide radicals, resulting in the generation of ONOO-, which increases cochlear nitrative stress, leading to the nitration of many cochlear proteins affecting their normal function. Several studies have reported that cisplatin treatment increases nitrotyrosine and ONOO^-^levels, which cause apoptosis in the cochlea, leading to hearing loss [48, 52]. Consistent with these reports, we observed a significant increase in nitrotyrosine levels in the hair cells and SGNs after cisplatin exposure.

Recent studies have provided potential insights into MnTBAP’s ability to alleviate the toxic side effects of cisplatin chemotherapy. For example, MnTBAP functioned as a mitochondrial ROS scavenger, efficiently decreasing mitochondrial ROS levels while restoring mitochondrial homeostasis in cisplatin-induced acute kidney injury models. It also mitigated oxidative and endoplasmic reticulum (ER) stress in cisplatin-induced nephrotoxicity [53]. In another study, MnTBAP cotreatment preserved cochlear synaptic integrity by reversing the cisplatin-induced decrease in cochlear synaptosomal proteins and attenuating cochlear synaptopathy [54]. Moreover, MnTBAP in its pure form acted as an effective scavenger of ONOO^-^ [45], a critical mediator of the ototoxic effects of cisplatin [48]. In this study, we observed a strong correlation between cisplatin-induced nitrative stress and hair cell and SGN loss. Because effective communication among hair cells, SGNs, and the auditory nerve is essential for normal auditory function [55, 56], and nitrative stress can promote the degradation of vulnerable nitrated proteins [57, 58], the attenuation of cisplatin-induced cochlear nitrative stress by MnTBAP cotreatment, along with the preservation of hair cells and SGNs, clearly demonstrates the otoprotective potential of MnTBAP.

The cisplatin-induced changes in the ABRs and DPOAEs were also consistent with the observed morphological changes in the hair cells and SGNs. Cisplatin treatment elevated hearing thresholds and caused a significant hearing threshold shift on day 8, which is consistent with the threshold shift observed with a cumulative dose of 15 mg/kg in previous reports [48, 59]. MnTBAP cotreatment attenuated the cisplatin-induced elevation in hearing thresholds, indicating its ability to preserve auditory signal transmission through the SGNs by protecting them from cisplatin-induced damage. Similarly, DPOAE amplitudes were significantly reduced following cisplatin treatment, whereas MnTBAP cotreatment mitigated this decrease, suggesting a protective effect on OHC function against cisplatin-induced damage. Overall, this study provides both morphological and physiological evidence supporting the therapeutic value of MnTBAP against cisplatin-induced ototoxicity.

## Conclusion

In summary, this study indicated that cotreatment with MnTBAP reversed the cisplatin-induced increase in cochlear nitrative stress, which is implicated in cochlear apoptosis, and protected the hair cells and SGNs from the toxic side effects of cisplatin. Additionally, the results demonstrated that MnTBAP cotreatment attenuated cisplatin-induced hearing loss and reduced the weight loss associated with cisplatin treatment. Collectively, these findings suggest that MnTBAP may serve as a promising therapeutic agent for mitigating cisplatin-induced ototoxicity.

## Acknowledgments

We acknowledge the assistance of the Wayne State University Microscopy, Imaging & Cytometry Resources Core.

## Author Contribution

Conceptualization: Samson Jamesdaniel; Methodology: Shomaila Mehmood, Pankaj Bhatia; Formal analysis and investigation: Shomaila Mehmood, Pankaj Bhatia; Writing - original draft preparation: Shomaila Mehmood; Writing - review and editing: Samson Jamesdaniel; Funding acquisition: Samson Jamesdaniel; Resources: Samson Jamesdaniel; Supervision: Samson Jamesdaniel.

